# ATR is a multifunctional regulator of male mouse meiosis

**DOI:** 10.1101/172106

**Authors:** Alexander Widger, Shantha K Mahadevaiah, Julian Lange, Elias Ellnati, Jasmin Zohren, Takayuki Hirota, Marcello Stanzione, Obah Ojarikre, Valdone Maciulyte, Dirk de Rooij, Attila Tóth, Scott Keeney, James MA Turner

## Abstract

Meiotic cells undergo genetic exchange between homologous chromosomes through programmed DNA double-strand break (DSB) formation, recombination and synapsis^1, 2^. In mice, the DNA damage-regulated phosphatidylinositol-3-kinase-like kinase (PIKK) ATM regulates all of these processes^3-6^. However, the meiotic functions of another major PIKK, ATR, have remained elusive, because germ line-specific depletion of this kinase is challenging. Using an efficient conditional strategy, we uncover roles for ATR in male mouse prophase I progression. Deletion of ATR causes chromosome axis fragmentation and germ cell elimination at mid pachynema. ATR is required for homologous synapsis, in a manner genetically dissociable from DSB formation. In addition, ATR regulates loading of recombinases RAD51 and DMC1 to DSBs and maintenance of recombination foci on synapsed and asynapsed chromosomes. Mid pachytene spermatocyte elimination in ATR deficient mice cannot be rescued by deletion of ATM and the third DNA damage-regulated PIKK, PRKDC, consistent with the existence of a PIKK-independent surveillance mechanism in the mammalian germ line. Our studies identify ATR as a multifunctional regulator of mouse meiosis.

---

ATR is a serine-threonine kinase with ubiquitous functions in somatic genome stability and checkpoint control^7^. Studies on non-mammalian model organisms have revealed that ATR is also essential for meiosis. ATR orthologues regulate meiotic DSB resection^8^, stoichiometry of DSB-associated strand-exchange proteins RAD51 and DMC1^9^, inter-homologue bias^10, 11^ and crossover formation^12^. They are also components of prophase I checkpoints that ensure centromere pairing^13^, timely repair of recombination intermediates^14, 15^, and correct coupling of DNA replication with DSB induction^16, 17^. Determining the functions of ATR in mouse meiosis has been hampered by the fact that ablation of this protein causes embryonic lethality^18, 19^. An inducible *Cre-ERT2* approach recently revealed that ATR regulates meiotic sex chromosome inactivation (MSCI), the silencing of the X and Y chromosomes in male meiosis, via serine-139 H2AFX phosphorylation (γH2AFX)^20^. However, this method resulted in partial rather than complete ATR depletion. We therefore sought a superior conditional strategy for dissecting additional meiotic ATR functions.

For this purpose, we generated male mice carrying one *Atr* floxed (*Atr^fl^*) allele, in which the exon 44 kinase domain of *Atr* is flanked by *loxP* sites^21^, and one *Atr* null (*Atr*^-^) allele, in which the first three coding exons of *Atr* are replaced by a neomycin selection cassette^19^. The resulting *Atr*^*fl*/-^ males also carried a transgene expressing *Cre* recombinase under the control of either a *Stra8* or *Ngn3* promoter fragment. *Stra8-Cre* is expressed from P3 (postnatal day 3)^22^, while *Ngn3-Cre* is expressed from P7^23, 24^ Testis weights at P30 were reduced three to fourfold in *Atr*^*fl*/-^ *Stra8-Cre* males and *Atr*^*fl*/-^ *Ngn3-Cre* males relative to *Atr*^*fl*/+^ *Cre* carrying controls, while body weights were unaffected (Fig. 1a). Western blotting showed that ATR protein was reduced in *Atr*^*fl*/-^ *Stra8-Cre* testes, and even more so in *Atr*^*fl*/-^ *Ngn3-Cre* testes (Fig. 1b). This finding supports previous evidence that the majority of testis ATR expression occurs in spermatocytes^20, 25^. Testis histology revealed germ cell failure at seminiferous tubule stage IV, corresponding to mid pachynema of meiosis, in both *Cre* models (Fig. 1c), reminiscent of findings in *Atr*^*fl*/-^ *Cre-ERT2* mice^20^. However, the stage IV elimination was clearly less robust in *Atr*^*fl*/-^ *Stra8-Cre* than *Atr*^*fl*/-^ *Ngn3-Cre* males, because in the former model elongating spermatids were observed in some testis sections (Fig. 1c inset). We therefore focused on *Atr*^*fl*/-^ *Ngn3-Cre* mice (hereafter *Atr*^*fl*/-^), with *Atr*^*fl*/+^ *Ngn3-Cre* (hereafter *Atr*^*fl*/+^) serving as controls.

**Figure 1:**
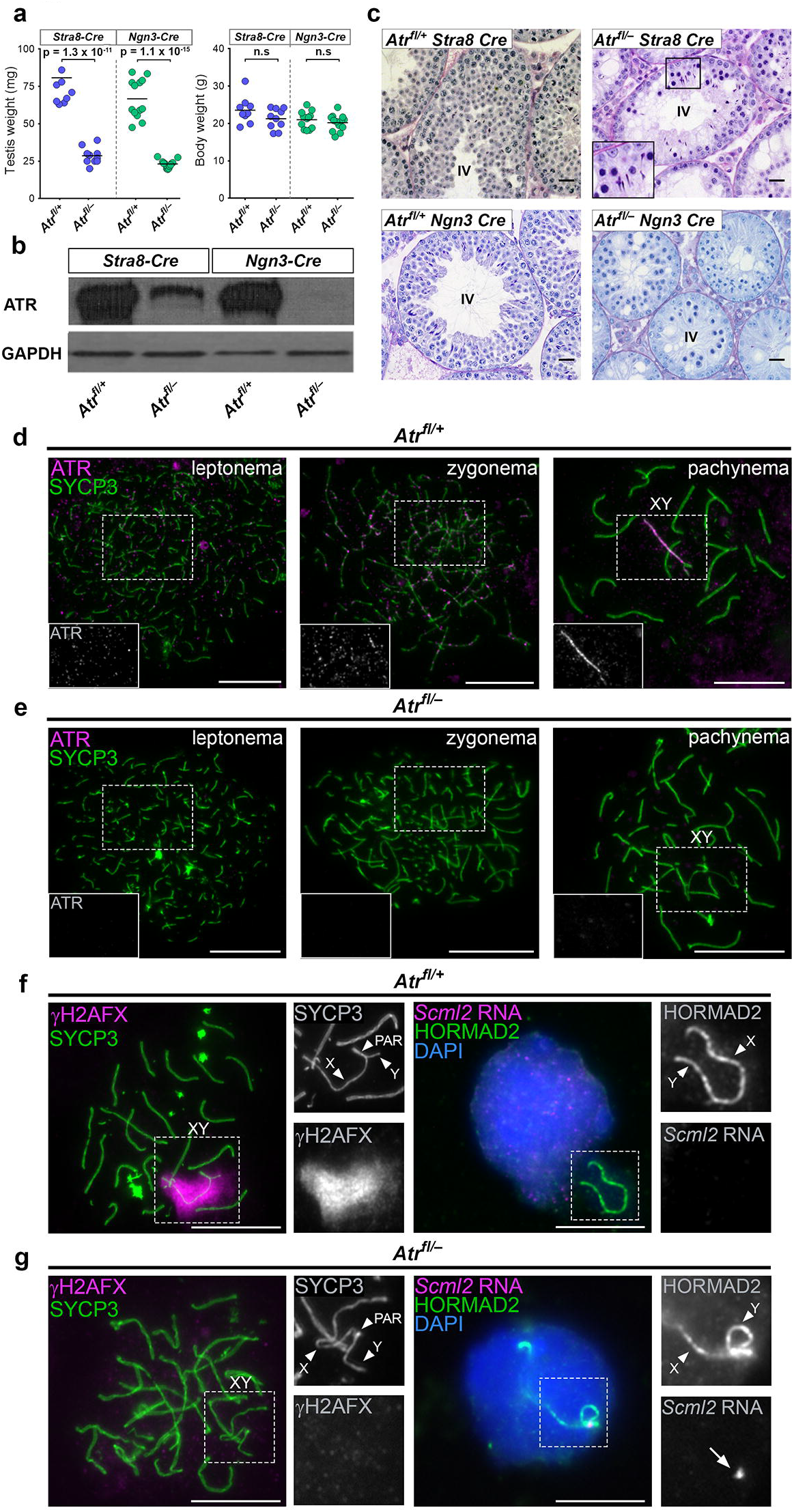
A conditional strategy for efficient depletion of ATR during male mouse meiosis. P30 testis and body weights (**a**), testis western blots (**b**), and Periodic Acid-Schiff and haemotoxylin / eosin stained stage IV testis sections (**c**) in *Atr*^*fl*/+^ *Stra8-Cre* males (n = 9 males), *Atr*^*fl*/-^ *Stra8-Cre* males (n = 10 males), *Atr*^*fl*/^+ *Ngn3-Cre* males (n = 13 males) and *Atr*^*fl*/-^ *Ngn3-Cre* males (n = 13 males; means and p values for **a** indicated; unpaired t test). Inset in (**c**) shows presence of elongating spermatids in some tubules from *Atr*^*fl*/-^ *Stra8-Cre* males. (**d,e**) ATR (magenta) and SYCP3 (green) staining in *Atr^fl/^+ Ngn3-Cre* (denoted Atr^fl/^+) and *Atr*^*fl*/-^ *Ngn3-Cre* (denoted *Atr^fl/-^*) males. In *Atr*^*fl*/+^ males (n = 2 males) ATR is observed as foci (see insets) in 85% of leptotene cells (n = 20 cells) and 82% of zygotene cells (n = 28 cells), and on the asynapsed region of the XY bivalent in 100% of pachytene cells (n = 30 cells). In *Atr*^*fl*/-^ males (n = 2 males) ATR is observed in no cells at these three stages (n = 21, 30 and 32 cells at leptonema, zygonema and pachynema, respectively). (**f**) Validation of ATR depletion in *Atr*^*fl*/-^ by analysis of MSCI. In *Atr*^*fl*/+^ males, all pachytene cells show coating of the X and Y chromosome (arrowheads) but not the pseuodautosomal regions (PAR, arrowhead) by γH2AFX (magenta; left panels; XY bivalent in box magnified in smaller panels). This coating causes silencing of the X chromosome gene *Scml2* (magenta; right panels) and compartmentalization of the XY bivalent (labeled with HORMAD2; green) in the sex body (n = one male, 29 cells; magnified in smaller panels). (**g**) In *Atr*^*fl*/-^ males, XY chromosome γH2AFX coating and sex body compartmentalization do not occur. As a result, *Scml2* expression (arrow) persists in all early pachytene cells (n = one male; 30 cells). Scale bar in (**c**) 20 μm, other scale bars 10 μm.

Combined immunofluorescence for ATR and the axial element protein SYCP3^26^ confirmed that the characteristic ATR staining pattern observed in control leptotene, zygotene and pachytene spermatocytes (Fig. 1d) was absent in *Atr*^*fl*/-^ males (Fig. 1e). Furthermore, MSCI, assayed at early pachynema by acquisition of γH2AFX on the XY bivalent and RNA FISH for expression of the X chromosome gene *Scml2*, was present in control males (Fig. 1f) but abolished in *Atr*^*fl*/-^ males (Fig. 1g). Thus, by multiple criteria, *Atr*^*fl*/-^ males exhibited efficient ATR depletion.

At stage IV, when wild type spermatocytes reach mid pachynema, *Atr*^*fl*/-^ spermatocytes contained highly fragmented chromosome axes and nucleus-wide γH2AFX staining (Supplementary Fig. 1a; see Methods for meiotic staging criteria used throughout this manuscript). These cells were readily distinguishable from *Atr*^*fl*/-^ cells at leptonema, in which axial elements were shorter and uniform in length, and γH2AFX staining across the nucleus was more heterogeneous (Supplementary Fig. 1b). Mid pachytene axis fragmentation and nucleus-wide γH2AFX staining were also noted in *Atm*^-/-^ males (Supplementary Fig. 1c), as described previously^27^. However, neither phenotype was observed in *Spo11*^-/-^, *Dmc1*^-/-^ and *Msh5*^-/-^ males (Supplementary Fig. 1d-f), which display stage IV arrest. Instead, γH2AFX in *Spo11*^-/-^ spermatocytes was restricted to the transcriptionally inactive pseudosex body (Supplementary Fig. 1d), while in *Dmc1*^-/-^ and *Msh5*^-/-^ spermatocytes it formed axis-associated clouds (Supplementary Fig. 1e,f), consistent with published reports^27–29^. These findings suggest that mid pachytene axis degeneration and nucleus-wide γH2AFX staining are features of ATR and ATM deletion, and not merely a consequence of stage IV germ cell death.

We then used *Atr*^*fl*/-^ males to address whether ATR regulates homologous synapsis. We performed immunostaining for SYCP3 and the asynapsis marker HORMAD2, focusing on *Atr*^*fl*/-^ cells at early pachynema, i.e. prior to extensive axial element fragmentation (Supplementary Fig. 1a). Synapsis was normal in 86% (n=104) of early pachytene cells from control males. As expected, HORMAD2 was absent on the autosomal bivalents and present on the non-homologous, asynapsed regions of the X and Y chromosome in these instances (Fig. 2a). Furthermore, in these cells, non-homologous synapsis between the sex chromosomes and the autosomes was not observed. However, in *Atr*^*fl*/-^ males, only 20% (n=107) of early pachytene cells achieved complete homologous synapsis. The remaining cells exhibited varying degrees of asynapsis affecting the XY pair and the autosomes (Fig. 2b; see below for details). In addition, non-homologous synapsis between the sex chromosomes and autosomes occurred in 25% of *Atr*^*fl*/-^ cells (Fig. 2b). In mice, the XY pseudoautosomal regions (PARs) undergo late synapsis and DSB formation^30^, and in yeast the homologue bias of late-forming DSBs is partially dependent on ATR^8^. We therefore determined whether asynapsis in *Atr*^*fl*/-^ males more commonly affects the sex chromosomes than the autosomes. To identify the sex chromosomes, we performed DNA FISH using probes for the X chromosome (*Slx* probe) and Y chromosome (*Sly* probe; Fig. 2a,b; insets). In *Atr*^*fl*/-^ males 77% of early pachytene cells exhibited XY asynapsis, while 59% exhibited autosomal asynapsis (see legend for further details). In control males, 13% of early pachytene cells exhibited XY asynapsis and 13% autosomal asynapsis. Thus ATR deletion has a more deleterious effect on XY than on autosomal synapsis.

**Figure 2:**
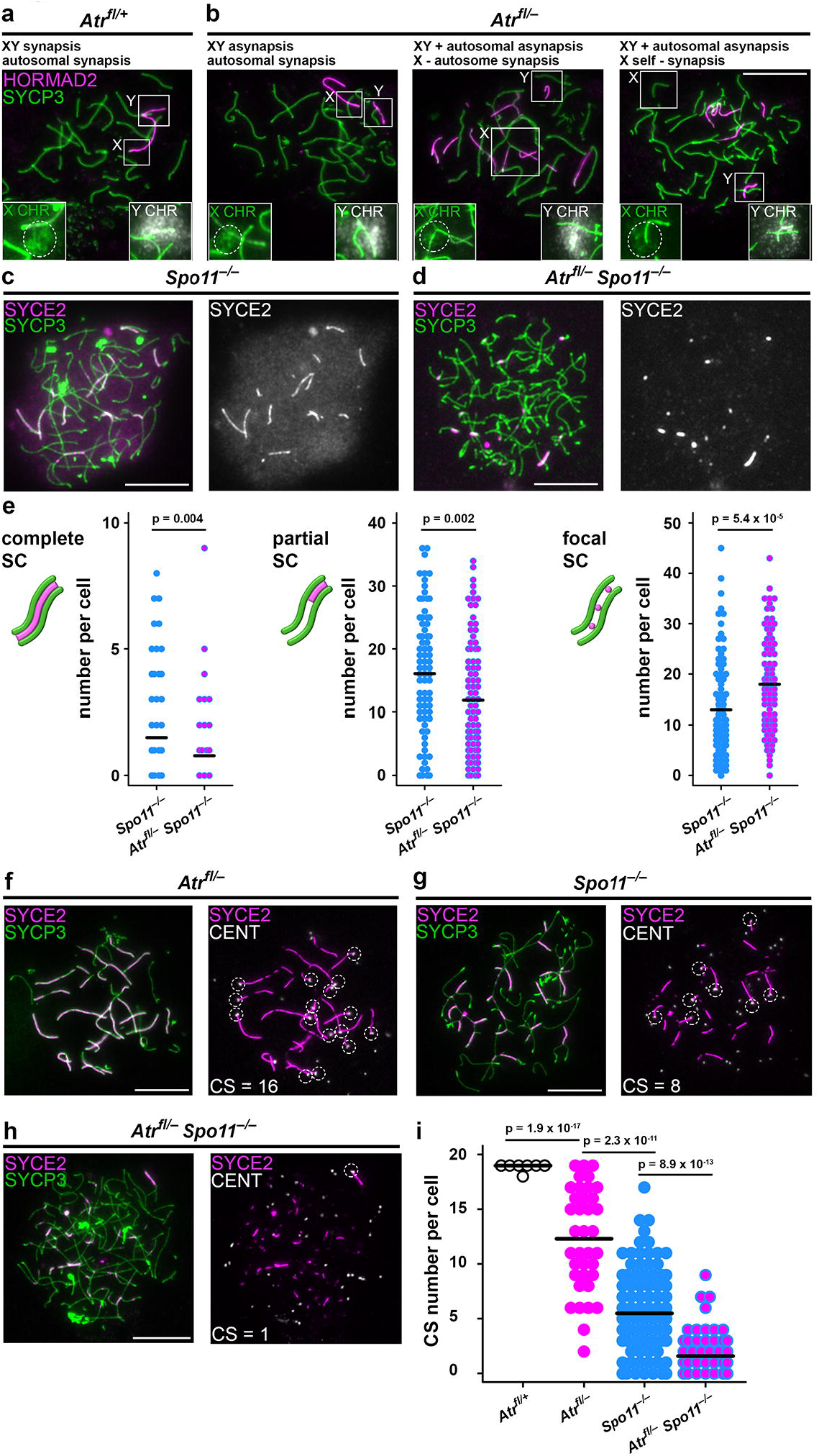
ATR is required for homologous synapsis. Examples of early pachytene synaptic outcomes in (**a**) *Atr*^*fl*/+^ males (n = 3 males) and (**b**) *Atr*^*fl*/-^ males (n = 3 males) assessed using HORMAD2 (magenta), SYCP3 (green) and subsequent DNA FISH using *Slx* and *Sly* probes (labeled in insets as X chromosome in green and Y chromosome in white). The *Slx* probe hybridizes to a sub-region of the X chromosome (circled), while the *Sly* probe coats the majority of the Y chromosome. In *Atr*^*fl*/+^ males (**a**) both the autosomes and the XY PARs are synapsed, while the non-homologous regions of the XY pair are asynapsed. In *Atr*^*fl*/+^ males (n=104 cells, 2 males), 90 cells had normal synapsis, 12 cells had asynapsis of both the XY and autosomes, 1 cell had asynapsis only of the XY and 1 cell had asynapsis only of the autosomes. (**b**) Three examples of synaptic defects in *Atr*^*fl*/-^ males, each described above respective image. In *Atr*^*fl*/-^ males (n=107 cells, 2 males), 21 cells had normal synapsis, 59 cells had asynapsis of both the XY and autosomes, 23 cells had asynapsis only of the XY and 4 cells had asynapsis only of the autosomes. (c,d) Comparison of pachytene synaptic outcomes in (**c**) *Spo11*^-/-^ and (**d**) *Atr*^*fl*/-^ *Spo11*^-/-^ males using immunostaining for SYCE2 (magenta) and SYCP3 (green). (**e**) Quantitation of complete, partial and focal SC in *Spo11*^-/-^ males (n = 2 males; 88 cells) and *Atr*^*fl*/-^ *Spo11*^-/-^ males (n = 2 males; 96 cells). Means and p values (unpaired t test) indicated. (f-h) Epistasis analysis of SPO11 and ATR in synapsis using the same markers plus immunostaining for centromeres (CENT; white) to determine CS number. For each cell, colocalizing SYCE2-CENT signals are indicated with dashed circles, and resulting CS numbers are shown. (**i**) CS number in *Atr*^*fl*/+^ males (n = 2 males; 55 cells), *Atr*^*fl*/-^ males (n = 2 males; 37 cells), *Spo11*^-/-^ males (n = 2 males; 114 cells) and *Atr*^*fl*/-^ *Spo11*^-/-^ males (n = 2 males; 64 cells). Mean values and p values (Mann-Whitney test) indicated. Scale bars 10 μm.

Asynapsis can result from defects in synaptonemal complex (SC) formation or recombination. To address whether ATR can promote SC formation independent of recombination, we used the *Spo11* null mutation, which permits genetic dissociation of synapsis from recombination initiation^31^. *Spo11*^-/-^ males do not form programmed DSBs, yet achieve extensive SC formation between non-homologues^32, 33^ If deletion of ATR impeded SC formation in *Spo11* nulls, then ATR must have a role in SC formation. We therefore compared synapsis between *Spo11*^-/-^ and *Atr^fl/-^ Spo11^-/-^* males. Like *Spo11*^-/-^ males, *Atr*^*fl*/-^ *Spo11*^-/-^ males exhibited stage IV germ cell elimination (Supplementary Fig. 2a,b). We classified SC formation, assessed using SYCP3 and the SC central element component SYCE2^34^, into three classes: i) complete SC, encompassing the entire axis length, ii) partial SC, extending along only part of an axis, and iii) focal SC (Fig. 2c-e). Relative to *Spo11*^-/-^ males (Fig. 2c), *Atr*^*fl*/-^ *Spo11*^-/-^ males (Fig. 2d) exhibited a decrease in complete and partial SC formation, and an increase in focal SC (Fig. 2c-e). These findings suggest that ATR promotes conversion of SC foci into longer SC stretches. Using *Atr^fl/-^ Spo11^-/-^* males, we were also able to demonstrate that formation of the pseudosex body in *Spo11*^-/-^ males is ATR-dependent (Supplementary Fig. 2c,d).

Our data suggested that ATR can promote synapsis independent of recombination. To strengthen these findings, we devised an additional method for higher throughput quantitation of synapsis across multiple genotypes. We triple immunostained cells for SYCP3, SYCE2 and centromeres, and counted the number of centromeres that had achieved synapsis, i.e. that colocalised with SYCE2 signals (centromere-SYCE2, or CS number; Fig. 2f-i). The mean CS number was reduced in *Atr*^*fl*/-^ relative to control males, confirming that ATR is required for normal levels of synapsis (Fig. 2f,i). The mean CS number was also lower in *Spo11*^-/-^ than in *Atr*^*fl*/-^ males, demonstrating that synapsis is more severely impaired by loss of SPO11 than loss of ATR (Fig. 2f,g,i). In *Atr*^*fl*/-^ *Spo11*^-/-^ males, the mean CS number was reduced relative to that in *Spo11*^-/-^ males (Fig. 2g,h,i). Thus, ATR promotes SC formation both in the presence and absence of SPO11-generated DSBs.

Next, we examined roles of ATR in DSB formation. Orthologues of ATM influence DSB homeostasis by acting as negative regulators of DSB induction^3, 35–38^. In *Saccharomyces cerevisiae* ATR promotes DSB formation indirectly by increasing the length of prophase I^39^, but its impact on DSB levels in mammals is unknown. To address this question, we measured abundance of covalent SPO11-oligonucleotide (SPO11 oligo) complexes, by-products of DSB induction, in testis extracts^40^. Consistent with previous work^3^, in *Atm*^-/-^ testes SPO11-oligo complex levels were elevated (Fig. 3a,b) and SPO11 oligos were shifted to a size larger than those observed in wild type mice (Fig. 3c,d). However SPO11-oligo levels and size distribution in *Atr*^*fl*/-^ testes were not detectably changed relative to controls (Fig. 3a-d). Thus, ATR and ATM have distinct functions with respect to DSB formation.

**Figure 3:**
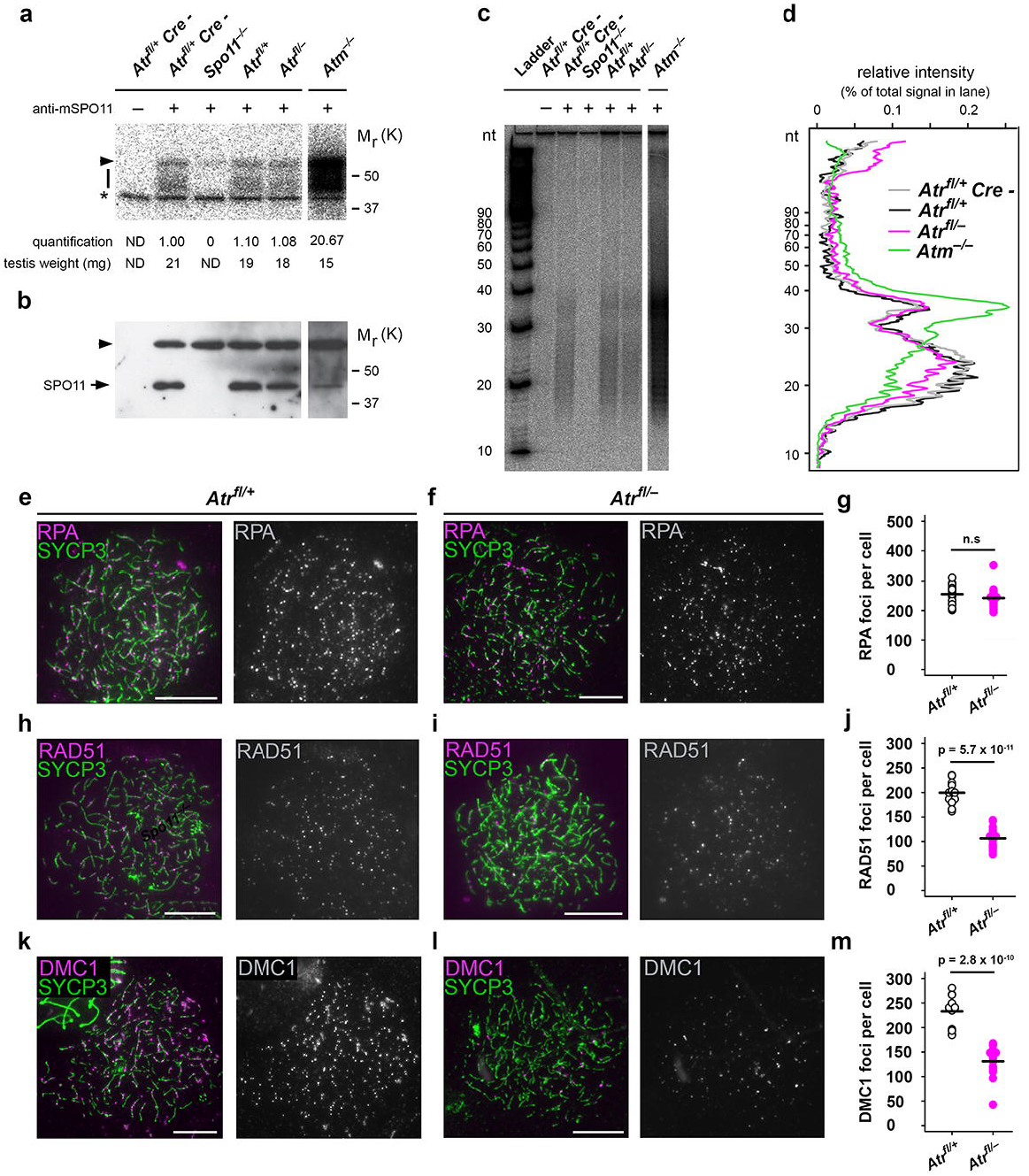
ATR ablation does not alter DSB levels but leads to reduction in leptotene recombinase counts. (**a-d**) Analysis of SPO11-oligo complexes in P13 testes. SPO11 was immunoprecipitated from whole-testis extracts and SPO11-associated oligos were end-labeled with terminal deoxynucleotidyl transferase and [α-^32^P] dCTP, then either separated on SDS-PAGE gels followed by autoradiography (**a**) and western blotting with anti-SPO11 antibody (**b**) or digested with proteinase K and resolved on denaturing polyacrylamide sequencing gels (c; background-subtracted lane traces in d). A representative experiment is shown. An additional *Atr*^*fl*/+^ *Cre* - control is shown to demonstrate that the *Ngn3-Cre* transgene does not influence SPO11-oligo levels. In (**a**) and (**b**), the bar indicates SPO11-oligo complexes, arrowhead indicates the immunoglobulin heavy chain, and asterisk marks non-specific labelling; ND = not determined. Each panel shows lanes from the same exposure of a single western blot or autoradiograph, with intervening lanes omitted. For quantitation, SPO11-oligo complex signals were background-subtracted and normalized to *Atr fl*/+ *Cre* - (n = 2) controls. SPO11-oligo quantitation: *Atr*^*fl*/+^ (1.05 ± 0.07-fold, mean and s.d., n = 2 males), *Atr*^*fl*/-^ (1.17 ± 0.22-fold, n = 4 males) and *Atm*^-/-^ (14.58 ± 5.24-fold, n = 3 males). (**e-m**) Analysis of leptotene focus counts using SYCP3 (green) and early recombination markers (magenta): RPA (**e-g**), RAD51 (**h-j**) and DMC1 (**k-m**) in *Atr*^*fl*/+^ males (n = 2 males; 27 cells for RPA, 19 cells for RAD51, 14 cells for DMC1) and *Atr*^*fl*/-^ males (n = 2 males; 21 cells for RPA, 19 cells for RAD51, 13 cells for DMC1). Mean and p values (Mann-Whitney test) indicated. Scale bars 10 μm.

During leptonema, resected DSBs are coated with RPA and the recombinases RAD51 and DMC1, which carry out strand invasion and recombinational repair. We used immunostaining to determine whether these early recombination components are influenced by ATR. Leptotene focus counts for RPA subunit 2 (hereafter termed RPA) were similar between *Atr*^*fl*/-^ and controls (Fig. 3e-g). This finding supports conclusions from SPO11-oligo complex quantitation that ATR does not influence

DSB abundance (Fig. 3a,b), and is consistent with RPA acting upstream of ATR^41^. Importantly though, in *Atr*^*fl*/-^ males, RAD51 counts were reduced by almost half relative to controls (Fig. 3h-j). DMC1 counts were lower by a similar magnitude (Fig. 3k-m). Thus, in *Atr*^*fl*/-^ males, DSB abundance appears grossly unaffected, but recombinase localization is compromised.

Next, we quantitated RPA, RAD51 and DMC1 foci during later prophase I. Interestingly, while RPA counts in *Atr*^*fl*/-^ males were equivalent to those in controls at late leptonema (Fig. 3e-g), they were reduced at mid zygonema (Fig. 4a-c). RAD51 and DMC1 counts were also lower in *Atr*^*fl*/-^ than control males at this stage (Fig. 4c; Supplementary Fig. 3a-d). At early pachynema, DSB markers were assayed both on the autosomes and on the X chromosome, focusing initially on cells with normal autosomal synapsis. Relative to control males, in *Atr*^*fl*/-^ males RPA, RAD51 and DMC1 counts were reduced on the autosomes (Fig. 4d-f; Supplementary Fig. 3e-h) and on the X chromosome (Fig. 4g,h,j; Supplementary Fig. 3i-l). The recombination defect in *Atr*^*fl*/-^ males therefore affects RAD51 and DMC1 at leptonema, and all three DSB markers at mid zygonema and early pachynema.

**Figure 4:**
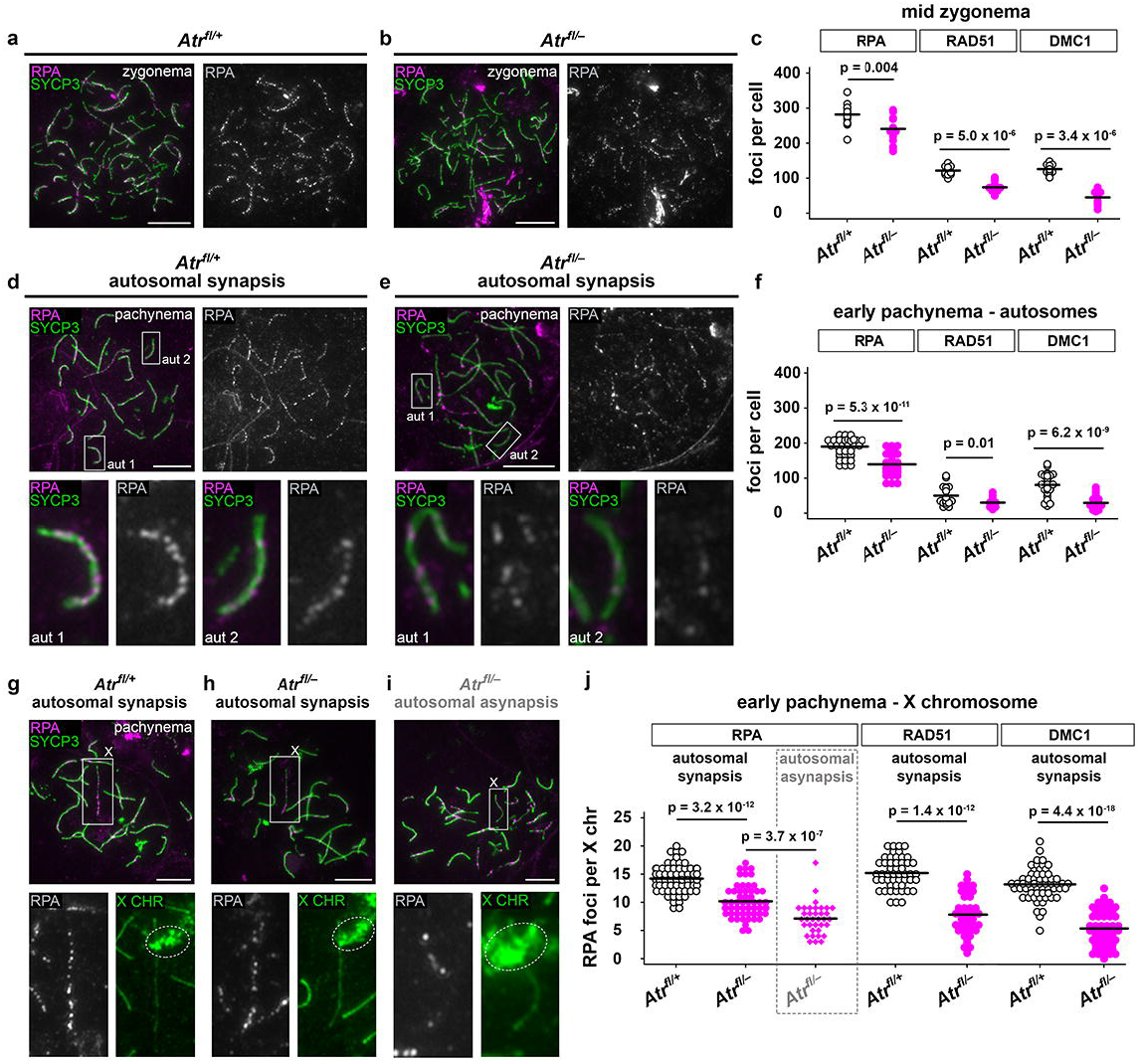
ATR regulates DSB marker counts during zygonema and pachynema. (**a,b**) RPA (magenta) and SYCP3 (green) immunostaining in *Atr*^*fl*/+^ and *Atr*^*fl*/-^ males. (**c**) RPA, RAD51 and DMC1 counts at mid zygonema in *Atr*^*fl*/+^ males (n = 2 males; 16 cells for RPA, 15 cells for RAD51, 15 cells for DMC1) and *Atr*^*fl*/-^ males (n = 2 males; 16 cells for RPA, 15 cells for RAD51, 15 cells for DMC1). (**d,e**) Examples of early pachytene cells from *Atr*^*fl*/+^ and *Atr*^*fl*/-^ males with normal autosomal synapsis. Two representative autosomes (aut 1 and 2) are boxed in upper panels and magnified in lower panels. (**f**) autosomal RPA, RAD51 and DMC1 counts at early pachynema in *Atr*^*fl*/+^ males (n = 2 males; 49 cells for RPA, 19 cells for RAD51, 33 cells for DMC1) and *Atr*^*fl*/-^ males (n = 2 males; 43 cells for RPA, 19 cells for RAD51, 35 cells for DMC1). (**g,h**) Early pachytene *Atr*^*fl*/+^ and *Atr*^*fl*/-^ cells with normal autosomal synapsis and the X chromosome (boxed in upper panels) identified by *Slx* DNA FISH (dashed circles in lower panels). (**i**) Early pachytene *Atr*^*fl*/-^ cell with autosomal asynapsis. (**j**) X chromosome RPA, RAD51 and DMC1 counts at early pachynema in *Atr*^*fl*/+^ males (n = 2 males; 64 cells for RPA, 49 cells for RAD51, 59 cells for DMC1) and *Atr*^*fl*/-^ males (n = 2 males; 62 cells for RPA in autosomal synapsis category, 35 cells for RPA in autosomal asynapsis category, 36 cells for RAD51, 53 cells for DMC1). Mean and p values (Mann-Whitney test) indicated. Scale bars 10 μm.

In mice, synapsis is dependent upon recombination. Asynapsis in *Atr*^*fl*/-^ males could therefore result not only from SC defects (Fig. 2), but also from aberrant recombination. We therefore asked whether early pachytene *Atr*^*fl*/-^ spermatocytes with autosomal asynapsis exhibit alterations in DSB marker counts relative to *Atr*^*fl*/-^ spermatocytes with normal autosomal synapsis. We used RPA as a representative DSB marker, and determined counts on the X chromosome as an indication of recombination intermediate levels. Since it can be obscured by asynapsed autosomes, the X chromosome was identified using *Slx* DNA FISH. In *Atr*^*fl*/-^ cells with autosomal asynapsis (Fig. 4i) X chromosome RPA counts were lower than those in *Atr*^*fl*/-^ cells exhibiting normal autosomal synapsis (Fig. 4h,j). Thus, greater alterations in recombination intermediates correlate with the asynapsis phenotype in *Atr*^*fl*/-^ males.

We then investigated mechanisms driving mid pachytene elimination of *Atr*^*fl*/-^ spermatocytes. In male mice, defective recombination causes germ cell loss via a mechanism promoted by ATM^42^. Since *Atr*^*fl*/-^ males exhibited a recombination defect (Fig. 3, 4), deletion of *Atm* in these mice might be expected to rescue their germ cell loss phenotype. However, *Atr*^*fl*/-^ males also exhibit defective MSCI, which can independently cause stage IV elimination^20, 43^ We therefore predicted that mid pachytene germ cell loss would be preserved in mice doubly deficient for ATR and ATM.

To distinguish these possibilities, we examined testis histology in *Atr*^*fl*/-^ *Atm*^-/-^ mutants. As expected, *Atr*^*fl*/-^ *Atm*^-/-^ testes exhibited efficient reduction in ATR and ATM protein levels (Fig. 5a,b). Leptotene and zygotene H2AFX phosphorylation, which are catalyzed by ATM and ATR, respectively^27, 28, 44, 45^, were also absent in *Atr*^*fl*/-^ *Atm*^-/-^ males, providing additional evidence that both PIKKs had been efficiently depleted in these double mutants (Fig. 5c, left and middle panel). Consistent with our prediction, stage IV elimination was preserved in *Atr*^*fl*/-^ *Atm*^-/-^ testes (Fig. 5d). Furthermore, the mid pachytene nucleus-wide γH2AFX staining observed in *Atr*^*fl*/-^ males (Supplementary Fig. 1a) and *Atm*^-/-^ males (Supplementary Fig. 1c) was also seen in *Atr*^*fl*/-^ *Atm*^-/-^ males (Fig. 5c, right panel).

**Figure 5:**
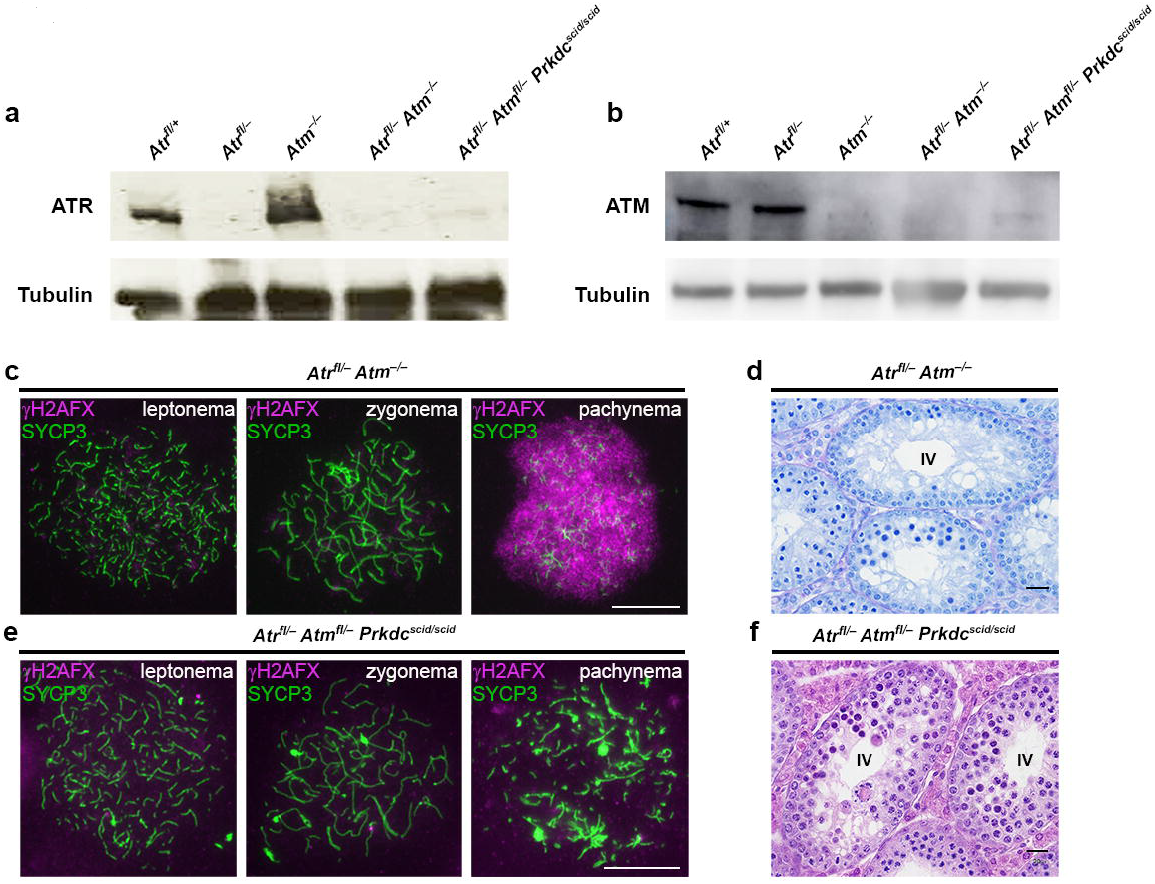
Mid pachytene germ cell elimination is preserved in mice deficient in the PIKKs. Western blot showing (**a**) ATR and (**b**) ATM depletion in mice with different PIKK mutations. (**c**) γH2AFX (magenta) and SYCP3 (green) immunostaining (n = 2 males, 25 cells for each stage) and (**d**) stage IV elimination in *Atr*^*fl*/-^ *Atm*^-/-^ males. (**e**) γH2AFX and SYCP3 immunostaining (n = 2 males, 25 cells for each stage) and (**f**) stage IV elimination in *Atr*^*fl*/-^ *Atm*^*fl*/-^ *Prkdc*^*scid/scid*^ males. Scale bar 20 μm in (**d,f**) and 10 μm in (**c,e**).

Our findings were consistent with a checkpoint-independent mechanism, most likely defective MSCI, driving germ cell loss in *Atr*^*fl*/-^ *Atm*^-/-^ males. However, the remaining DNA damage-regulated PIKK, *Prkdc*, was still present in these mutants, and could contribute a checkpoint function in the absence of ATR or ATM. We therefore examined germ cell progression in males deficient in all three DNA damage-regulated PIKKs. For this experiment we used the *Prkdc scid* mutation^46^ and an *Atm flox* allele^47^, because combined homozygosity for *Prkdc scid* and the *Atm* null mutation causes embryonic lethality^48^. In the resulting *Atr*^*fl*/-^ *Atm*^*fl*/-^ *Prkdc*^*scid/scid*^ males, testis ATM and ATR levels were depleted (Fig. 5a,b), and consequently ATM and ATR-dependent γH2AFX staining was absent (Fig. 5e, left and middle panels). Interestingly, mid pachytene nucleus-wide γH2AFX staining was also abolished (Fig. 5e, right panel). Thus, PRKDC mediates pachytene serine-139 H2AFX phosphorylation in *Atr*^*fl*/-^ *Atm*^*fl*/-^ males. Nevertheless, in *Atr*^*fl*/-^ *Atm*^*fl*/-^ *Prkdc*^*scid/scid*^ testes stage IV germ cell loss still occurred (Fig. 5f). Thus, mid pachytene elimination persists in mice deficient in all three DNA damage-regulated PIKKs.

The functions of ATR in mammalian meiosis have been unclear. We show here that ATR is required during unperturbed meiosis to regulate chromosome axis integrity, synapsis and recombination. Notably, as is the case in mitosis^7^, in meiosis ATR has roles that are both shared and distinct from ATM. Both ATM and ATR are required for synapsis, but they exert different influences on DSB levels and on recombination focus constituents. Such differences are likely explained by the contrasting substrate Specificities^7^ and meiotic expression profiles^28^ of these kinases.

Our findings show that ATR can promote synapsis independently of meiotic DSB formation, presumably through modification of SC proteins. Among the many SC components, HORMAD1/2 and SMC3 are established ATR phosphortargets^20, 49, 50^. While additional SC candidates no doubt exist^51, 52^, HORMAD1 is of particular interest, because this protein can also promote synapsis in the absence of recombination^31^. We find that in wild type males, asynapsis of the XY pair occurs at similar frequency to that of all autosomal pairs combined. This bias towards XY asynapsis may be attributable to the small length and terminal location of the PAR. Notably however, in *Atr* mutants, the bias is exaggerated, such that asynapsis of the XY bivalent occurs at higher frequency than that of all autosomes combined. The PAR is unusual, undergoing late DSB formation and synapsis^30^, and being enriched for repressive chromatin marks not observed at the termini of autosomal bivalents^53, 54^ We suggest that ATR promotes one or more of these unique properties to ensure successful XY interactions.

At leptonema in ATR-deficient males, RAD51 and DMC1 counts are reduced, while RPA counts are unaffected. This finding could indicate a role for ATR in RAD51/DMC1 loading. It has been suggested that ATR phosphorylates RAD51^55^. ATR also phosphorylates CHK1 during meiosis^56^, and CHK1 in turn promotes RAD51 loading at DSBs^57^. Subsequently, at zygonema and pachynema, spermatocytes lacking ATR exhibit lower counts not only for RAD51/DMC1 but also for RPA. This reduction in all DSB markers could imply additional functions for ATR in later processing of recombination intermediates. Alternatively, it may indicate the presence of fewer unrepaired DSBs. Reduced DSB number could, in turn, result from premature repair using the sister chromatid^11^, or from a failure to induce new DSBs on asynapsed chromosomes^58^. Interestingly, the ATR substrates HORMAD1/2 are implicated both in inhibiting sister chromatid recombination^59, 60^ and in promoting ongoing DSB formation^59, 61^. Our data are less consistent with a defect in generating new DSBs on asynapsed chromosomes, because SPO11-oligo data showed no reduction in DSB formation in ATR mutants. Furthermore, in ATR mutants decreased RPA counts were also observed on synapsed autosomes, and the magnitude of the decrease was similar to that seen for the asynapsed X chromosome. Whether ATR functions in later stages of recombination, e.g. crossover formation, is unclear, since late recombination markers appear after the point of arrest in *Atr*^*fl*/-^ germ cells (Supplementary Fig. 4).

An interesting finding is that mid pachytene elimination persists in mice deficient in all three DNA damage-regulated PIKKs. Multiple lines of evidence suggest that PIKK depletion in these mice is efficient; nevertheless the use of conditional alleles means that residual PIKK activity may be present and sufficient for checkpoint maintenance. Setting aside this caveat, our findings do not exclude a contribution of PIKKs to mid pachytene elimination in synapsis and recombination mutants. However, they do confirm the existence of additional mechanisms that can trigger elimination. Under such circumstances, mid pachytene failure is likely caused by defective MSCI^43^. We suggest that the coexistence of multiple overlapping surveillance mechanisms during prophase I in males explains why checkpoint responses are more robust than those in females^62, 63^. Further insight into this important sexual dimorphism will require the application of alternative conditional strategies for PIKK deletion in the female mouse germ line.

## Methods

### Animal experiments

All mice were maintained according to UK Home Office Regulations at the National Institute for Medical Research (NIMR) and the Francis Crick Institute Mill Hill laboratory. Genetically modified models are previously published and are maintained on a predominantly C57BL/6 background: *Ngn3-Cre^24^, Stra8-Cre^22^, Atr^19^, Atr fiox^64^, Atm^65^, Atm flox^47^, Spo11^33^, Dmc1^66^. Prkdc scid/scid* mice were obtained from the Jackson Labs and are maintained on an NOD background. Littermate controls were used where possible.

### Immunofluorescence and western blotting

Immunofluorescence experiments on surface spread spermatocytes were carried out as described^44^ In brief, cells were permeabilized for 10 min in 0.05 % Triton X-100 and fixed for one hour min in 2 % formaldehyde, 0.02 % SDS in phosphate-buffered saline (PBS). Slides were rinsed in distilled water, air dried and blocked in PBT (0.15 % BSA, 0.10 % TWEEN-20 in PBS) for one hour. Slides were incubated with the following antibodies in a humid chamber overnight at 37°C: guinea-pig anti-SYCP3 (made in-house) 1:500, rabbit anti-SYCP3 (Abcam ab-15092) 1:100, rabbit anti-ATR (Cell Signalling #2790) 1:50, mouse anti-γH2AFX (Millipore 05-636) 1:100, rabbit anti-HORMAD2 (Tóth lab) 1:100, guinea-pig anti-SYCE2 (gift from Howard Cooke / Ian Adams) 1:800, human anti-centromere (CREST ab gift from Bill Earnshaw) 1:1000, rabbit anti-RPA (Abcam ab-2175) 1:100 anti-rabbit RAD51 (Calbiochem PC130) 1:25, goat anti-DMC1 (Santa Cruz sc-8973) 1:25. Western blotting was carried out as described^67^.

### Meiotic staging

The presence of asynapsis can make discrimination between zygotene cells and pachytene cells with asynapsis challenging. We used the following criteria: (1) DAPI staining: at zygonema DAPI staining is bright and centromeres are clustered in a few subdomains, while in pachynema DAPI staining is heterogeneous, with euchromatin stained faintly and centromeres stained brightly and forming multiple subdomains. (2) The length and thickness of axial elements / SCs: axial elements are long and thin in zygonema, but shorter and thicker in pachynema. (3) Chromosomal asynchrony: in contrast to zygotene cells, pachytene cells with asynapsis exhibit asynchrony in synapsis between individual bivalents, i.e. the coexistence of completely asynapsed bivalents and multiple fully synapsed bivalents. We determined that *Atr*^*fl*/-^ males cells exhibiting chromosome axis fragmentation were at mid pachytene by analysis of SYCP3 staining in testis sections; where stage IV tubules can be readily identified. For comparison of synapsis between *Spo11*^-/-^ males and *Atr*^*fl*/-^ *Spo11*^-/-^ males, pachytene cells were identified by virtue of having fully developed axial elements.

### RNA, DNA FISH and SPO11-oligo analysis

FISH was carried out with digoxigenin-labeled probes as previously described^68^. CHORI BAC probe, RP24-204O18 (CHORI) was used for *Scml2* RNA FISH, RP23-470D15 for *Slx* DNA FISH, and RP24-502P5 for *Sly* DNA FISH. Analyses of abundance of SPO11-oligo complexes and sizes of SPO11 oligos were performed as described^3, 40^.

### Microscopy

Imaging was performed using an Olympus IX70 inverted microscope with a 100-W mercury arc lamp. For chromosome spread and RNA FISH imaging, an Olympus UPlanApo 100x/1.35 NA oil immersion objective was used. For testis section imaging, an Olympus UPlanApo 40x/0.75 NA objective was used. A Deltavision RT computer-assisted Photometrics CoolsnapHQ CCD camera with an ICX285 Progressive scan CCD image sensor was utilized for image capture. 8 or 16-bit (512x512 or 1024x1024 pixels) raw images of each channel were captured and later processed using Fiji software.

### Statistics

Statistical calculations were performed using GraphPad Prism 6.0. For comparison of two genotypes, Mann-Whitney test or t-tests were performed.

## Acknowledgements

We are grateful to Bill Earnshaw for providing the CREST anti-centromere antibody Howard Cooke, Ian Adams for providing the SYCE2 antibody, Paula Cohen for providing the anti-MLH3 antibody, Eric Brown for providing the *Atr* flox and null mice, members of the Turner laboratory for critical reading of the manuscript, and the Francis Crick institute technology platforms for excellent assistance. This work was supported by the Francis Crick Institute which receives its core funding from Cancer Research UK (FC001193), the UK Medical Research Council (FC001193), and the Wellcome Trust (FC001193). J.L. was supported in part by US National Institutes of Health grant R01 GM105421 (to Maria Jasin and S.K.) and by American Cancer Society fellowship PF-12-157-01-DMC.

## Author contributions

A.W, O.O, V.M and J.M.A.T performed animal generation and genotyping, A.W, S.K.M, M.S and J.M.A.T performed immunofluorescence, A.W, E.E and V.M performed western blots, A.W and S.K.M performed RNA FISH, T.H performed DNA FISH, J.L and S.K performed SPO11-oligo blots, A.W. and D. de. R performed histology, A.W and J.Z performed data plotting and statistics, A.T supplied HORMAD2 antibody. J.T wrote the manuscript with critical input from A.T, S.K and Turner lab members.

